# Reemergence of the Murine Bacterial Pathogen *Chlamydia muridarum* in Laboratory Mouse Colonies

**DOI:** 10.1101/2022.05.16.491822

**Authors:** Noah Mishkin, Rodolfo J Ricart Arbona, Sebastian E Carrasco, Samira Lawton, Kenneth S. Henderson, Panagiota Momtsios, Ira M Sigar, Kyle H Ramsey, Christopher Cheleuitte-Nieves, Sebastien Monette, Neil S Lipman

## Abstract

*Chlamydia muridarum* (Cm) was detected in mice from 2 colonies with lymphoplasmacytic pulmonary infiltrates using PCR and immunohistochemistry. This discovery was unexpected as Cm infection had not been reported in laboratory mice since the 1940’s. A Cm specific PCR assay was developed and testing implemented for resident colonies from 8 vivaria from 3 academic institutions, 58 incoming mouse shipments from 39 academic institutions, and mice received from 55 commercial breeding colonies (4 vendors). To estimate Cm’s global prevalence in laboratory colonies, a database containing 11,387 metagenomic fecal microbiota samples from 120 institutions and a cohort of 900 diagnostic samples from 96 institutions were examined. Results indicate significant prevalence amongst academic institutions with Cm detected in 62.9% of soiled bedding sentinels from 3 institutions; 32.7% of incoming mouse shipments from 39 academic institutions; 14.2% of 120 institutions submitting microbiota samples; and 16.2% of the diagnostic sample cohort. All samples from commercial breeding colonies were negative. Additionally, naïve NOD.Cg-*Prkdc*^scid^ Il2rg^tm1Wjl^/SzJ (NSG) mice exposed to Cm shedding mice and their soiled bedding developed clinical disease 21 to 28 days following exposure. These mice had a moderate-to-severe histiocytic and neutrophilic bronchointerstitial pneumonia with respiratory epithelium demonstrating inclusions, chlamydial major outer membrane protein immunostaining, and hybridization with a Cm reference sequence (GenBank accession no. U68436). Cm was isolated on HeLa 229 cells from lungs, cecum, and feces of a Cm infected NSG mouse. The considerable prevalence of Cm is likely attributed to widespread global interinstitutional distribution of unique mouse strains and failure to recognize that some of these mice were from enzootically infected colonies. Given that experimental Cm colonization of mice results in a robust immune response and, on occasion, pathology, natural infection may confound experimental results. Therefore, Cm should be excluded and eradicated from endemically infected laboratory mouse colonies.

## Introduction

Chlamydiae are obligate intracellular bacteria with an extensive host range at the genus level and high host specificity at the species level.^12,31^ *Chlamydia muridarum* (Cm) is the only natural chlamydial pathogen of mice identified to date. It has been used extensively to model sexually transmitted *Chlamydia trachomatis* infection of humans.^12,31^ Cm was first described in the late 1930’s and further investigated in the early 1940’s when scientists identified a putative infectious organism causing respiratory disease and lung pathology in mice used in studies of influenza and “the virus of the common cold”.^11,16^ The etiologic agent was initially referred to as “mouse pneumonitis virus (MoPn)” by Clara Nigg in 1944.^25^ It was subsequently identified as a biovar of *C. trachomatis* and finally as Cm after phylogenetic analysis, genome sequencing, and comparative genomics.^7,13,31,32,33^ Despite widespread use as an experimental model, relatively little is known about the biology of natural infection as Cm has not been routinely reported in or isolated from laboratory mouse populations since its initial discovery. A closely related strain (denoted as strain SFPD) was isolated from hamsters in the early 1990’s.^13,36^ This strain was shown to share 91% sequence homology of the major outer membrane (MOMP) with the Weiss and Nigg strains of Cm.^43^ While Cm was likely prevalent in early 20^th^ C laboratory mouse colonies, the development and introduction of modern biosecurity practices, e.g., C-section rederivation and barrier husbandry, in the middle of the 20^th^ C would have been expected to eliminate the bacterium, if present, from most vendors’ colonies. Accordingly, testing of commercial and research colonies is not routinely performed.

As with other chlamydiae, Cm has a biphasic life cycle comprised of the non-replicating but infectious “elementary body” (EB), and the replicating but non-infectious “reticulate body” (RB).^12,31^ EBs enter host mucosal epithelial cells and are incorporated into a membrane-bound compartment (also called an “inclusion” or “inclusion body,” IB) in which they differentiate into RB. ^12,31^ RBs replicate within inclusions eventually re-differentiating into EBs, whereby they are released and infect nearby cells.^12,31^ Natural transmission is now thought to occur primarily via the fecal-oral route, similar to chlamydial infections in domestic animals where the gastrointestinal tract is often the natural site of colonization.^42^ This is supported by experimental infection, as orally inoculated immunocompetent mice remain persistently infected, shedding for up to 260 days with replication primarily in the cecum and colon. Pulmonary lesions and colonization, as reported in the initial discovery of Cm and in subsequent experimental infections, are presumed to occur through aspiration or inhalation of the bacterium.^13,25,31,42^ There are significant differences in host response to Cm infection dependent on the mouse strain inoculated, with Cm’s MOMP considered an important antigenic stimulus.^22,31^

Following the surprising identification of Cm in association with lymphoplasmacytic pulmonary infiltrates in 2 genetically engineered mouse (GEM) strains from distinct colonies, a multifaceted investigation was undertaken to determine the prevalence of Cm within our and other institutional mouse colonies, as well as commercial vendors following development of a Cm specific PCR assay. We further confirmed the global prevalence of Cm in research mouse colonies by examining fecal microbiome data from a large dataset and testing a large cohort of multi-institutional diagnostic samples and successfully isolated Cm in cell culture from highly immunocompromised mice used as contact and soiled bedding sentinels.

## Materials and Methods

### History/Index Cases

In 2020, Cm was initially detected in 2 unrelated GEM strains maintained at our institution by 2 laboratories. Laboratory A reported unexpected inflammatory infiltrate on histology of the lungs in an experimentally naïve, immunocompetent, clinically normal GEM strain. Histopathology of the lungs revealed mild to moderate, multifocal, perivascular lymphocytic aggregates in 1 of 3, 6-month-old Foxp3^DTRγ^CD4^CreERT2^R26^tdTomato^, knock-in reporter mice on a C57BL/6 background submitted for evaluation. While there was no histologic evidence of bacterial or fungal pathogens, agents known to colonize the respiratory tract and cause inflammation are not always detectable histologically. Accordingly, formalin-fixed paraffin embedded (FFPE) scrolls of lung tissue were assessed for *Mycoplasma spp*., *Pneumocystis spp*., and Cm by real time PCR (Molecular Diagnostics, Auburn University, Auburn, AL). Polymerase chain reaction targeting of the highly conserved chlamydial 23S rRNA gene with duplex fluorescent oligonucleotide probes followed by sequencing of the resulting amplicon surprisingly confirmed the presence of Cm. ^10^ *Mycoplasma* spp. and *Pneumocystis* spp. were not detected via commercial PCR assays (IDEXX Bioanalytics, Columbia, MO). Two additional mice (C57Bl/6-Foxp3^tm1Flv^/J; C57Bl/6/Foxp3^CreER^R26^tdTomato^) were submitted from this laboratory’s colony in 2021 for further analysis including complete blood count, serum biochemistry, histopathology, immunohistochemistry (IHC) for MOMP, and in-situ hybridization (ISH) for Cm mRNA (see “Results” for details).

Subsequently, complete necropsies were conducted on 2, 4-week-old, female Cd23-Cre Gen1^-/-^ Mus81^fl/fl^ (conditional knockouts of DNA repair proteins in B cells on a C57BL/6 background) from Laboratory B’s rodent colony. These animals initially presented for runting, mild hunched posture, and alopecia and thinning of hair along the dorsum and bilateral flanks. Histology of the skin was characterized by follicular dystrophy compatible with a background lesion described in C57BL/6 mice.^39^ Both mice also had mild, multifocal, chronic lymphoplasmacytic bronchiolitis and bronchitis. Given the pulmonary lesions noted and the discoveries detailed for Laboratory A, additional diagnostics were pursued including MOMP IHC, ISH for Cm mRNA in lung tissue, and PCR (see “Results” for details).

### Investigative Plan

Following the unexpected detection of Cm in mice from 2 distinct institutional GEM colonies, a multifaceted investigation was undertaken to determine whether this finding was unique and, if Cm was enzootic in other colonies, how widely it was distributed.

After developing a Cm specific PCR assay (described below), the following investigations were initiated: 1) All incoming GEM strains (n=58) from non-commercial colonies were tested for Cm by fecal PCR; 2) Soiled bedding SW:Tac sentinels employed in our multi-institutional colony health monitoring program (described below) from 97 mouse holding rooms were tested by fecal PCR during their annual testing submissions; 3) Two NOD.Cg-*Prkdc*^scid^ Il2rg^tm1Wjl^/SzJ (NSG) mice were co-housed with 2 cages containing 4 mice each from a recently imported Cm fecal PCR-positive, immunocompetent, GEM strain with a floxed tumor-suppressor allele from another academic institution to increase the likelihood of isolating Cm. After cohousing as contact sentinels for 1 week, each of these NSG mice were subsequently cohoused with 4 naïve NSG mice in 2 cages which continued to receive soiled bedding from the imported mice for an additional week. The mice were observed until morbidity developed, at which point they were euthanized, a complete necropsy undertaken, IHC for MOMP and ISH was performed on select tissues, and samples of lung, cecum, and feces from a single morbid NSG contact sentinel were collected for subsequent isolation of Cm. The PCR product obtained from FFPE pulmonary scrolls from a single NSG sentinel mouse was sequenced and compared to published Cm sequences; 4) Cm PCR was performed on feces collected on arrival from mice sourced from 55 production rooms from 4 commercial breeding colonies. Additionally, several non-laboratory mouse populations were surveyed to determine whether Cm was present in mice outside of the laboratory environment. Feces were collected from *Mus musculus* from 3 pet stores (pet shop A and B in NYC, pet shop C in Michigan), and from a wild population of *Mus musculus* in New Jersey; 5) Anonymized diagnostic samples (n=900) which included feces; oral or body/fur swabs; tissue (mediastinal lymph node, lung, or spleen); environmental (IVC exhaust air dust, plenum/filter/hose/environmental swabs, or filter material); and, biologics (cells and/or tissue culture) were tested for Cm by PCR; and, 6) A database containing mouse fecal metagenomic microbiome sequencing data from 11,387 samples from global academic (n=112) and commercial (n=8) sources was queried for Cm abundance.

### Animals

Swiss Webster (Tac:SW; Taconic Biosciences, Germantown, NY) and NOD.Cg-*Prkdc*^*scid*^ *Il2rg*^*tm1Wjl*^/SzJ (NSG; The Jackson Laboratory, Bar Harbor, ME) were used. All mice were free of mouse hepatitis virus (MHV), Sendai virus, mouse parvovirus (MPV), minute virus of mice (MVM), murine norovirus (MNV), murine astrovirus 2 (MuAstV2), pneumonia virus of mice (PVM), Theiler meningoencephalitis virus (TMEV), epizootic diarrhea of infant mice (mouse rotavirus, EDIM), ectromelia virus, reovirus type 3, lymphocytic choriomeningitis virus (LCMV), K virus, mouse adenovirus 1 and 2 (MAD 1/2), polyoma virus, murine cytomegalovirus (MCMV), mouse thymic virus (MTV), Hantaan virus, mouse kidney parvovirus (MKPV), *Mycoplasma pulmonis, CAR bacillus, Chlamydia muridarum, Citrobacter rodentium, Rodentibacter pneumotropicus, Helicobacter spp*., segmented filamentous bacterium (SFB), *Salmonella* spp., *Streptococcus pneumoniae, Beta-hemolytic Streptococcus spp*., *Streptobacillus moniliformis, Filobacterium rodentium, Clostridium piliforme, Corynebacterium bovis, Corynebacterium kutscheri, Staphylococcus aureus, Klebsiella pneumoniae, Klebsiella oxytoca*,, *Pseudomonas aeruginosa*, fur mites (*Myobia musculi, Myocoptes musculinis*, and *Radfordia affinis*), pinworms (*Syphacia* spp. and *Aspiculuris* spp.), *Demodex musculi, Pneumocystis spp*,, *Giardia muris, Spironucleus muris, Entamoeba muris, Tritrichomonas muris*, and *Encephalitozoon cuniculi* when the studies were initiated.

### Husbandry and Housing

Mice were maintained in individually ventilated, polysulfone shoebox cages with stainless-steel wire-bar lids and filter tops (experimental and colony; no. 19, Thoren Caging Systems, Inc., Hazelton, PA) or in solid-bottom polysulfone microisolator cages maintained in a custom-designed quarantine caging system (non-commercial mice imported from other institutions were housed in a dedicated quarantine facility; BioZone Global, Chester, SC) on autoclaved aspen chip bedding (PWI Industries, Quebec, Canada) at a density of no greater than 5 mice per cage. Each cage was provided with a Glatfelter paper bag containing 6 g of crinkled paper strips (EnviroPak^®^, WF Fisher and Son, Branchburg, NJ) for enrichment. Mice were fed either a natural ingredient, closed source, flash-autoclaved, γ-irradiated feed (experimental and colony; LabDiet 5053, PMI, St. Louis, MO) or γ-irradiated feed containing 150 PPM fenbendazole and 13 PPM ivermectin (quarantine; TestDiet, PMI) *ad libitum*.^40^ All animals were provided autoclaved reverse osmosis acidified (pH 2.5 to 2.8 with hydrochloric acid) water in polyphenylsulfone bottles with stainless-steel caps and sipper tubes (Techniplast, West Chester, PA) ad libitum. Cages were changed every 7 days within a class II, type A2 biological safety cabinet (LabGard S602-500, Nuaire, Plymouth, MN) or an animal changing station (Nuaire NU-S612-400, Nuaire, Plymouth MN). The rooms were maintained on a 12:12-h light:dark cycle, relative humidity of 30% to 70%, and room temperature of 72 ± 2°F (22.2 ± 1.1°C). Rooms were classified as receiving animals from all sources, (e.g., non-commercial sources [released after testing for specific agents] and commercial breeders) versus from non-commercial sources (rederived into axenic state and reassociated with defined, vendor-specific flora) and/or commercial breeders only. The animal care and use program at Memorial Sloan Kettering Cancer Center (MSK) is accredited by AAALAC International, and all animals are maintained in accordance with the recommendations provided in the Guide.^30^ All animal use described in this investigation was approved by MSK’s IACUC in agreement with AALAS’ position statements on the Humane Care and Use of Laboratory Animals and Alleviating Pain and Distress in Laboratory Animals.

### Mouse Importation Programs and Colony Health Monitoring

Mouse importations from non-commercial sources, e.g., academic institutions and atypical commercial sources (commercial research companies, non-approved commercial breeders, and approved commercial breeders who perform insufficient testing) are subject to quarantine and testing. The length of quarantine and scope of testing are determined by several factors including intended use and health status. No imported mice were tested for Cm prior to arrival as institutions and companies are not regularly testing for this agent. Transport crates are received into the quarantine facility, where they are sprayed with chlorine dioxide (Clidox-S 1:18:1 dilution, Pharmacal Laboratories, Naugatuck, CT) and placed in the assigned holding room in a dedicated, physically separated, quarantine facility operated at ABSL-2. Testing is comprised of at least 2 rounds of testing and involves either all mice (when 10 or fewer are received), or a minimum of 10 mice with at least 1 from each cage received. Initial testing (QA) is performed within 72 hours of arrival; this testing includes veterinary assessment and PCR on fecal pellet, oral and fur/pelt swabs. Secondary testing (QB) is performed 4 weeks after QA, and also includes serology (with 0.25 mL blood collected via the sub-mandibular vein from 1 animal per cage). PCR and serologic samples are submitted to a commercial diagnostic laboratory (Charles River Laboratories, Cambridge, MA) for detection of the above listed agents (see “Animals”). Testing for the presence of Cm was added to the routine quarantine testing panel in October 2021.

The soiled bedding sentinel colony health monitoring program has been previously described.^20^ In brief, 3-5 week old female SW:Tac mice are utilized as soiled bedding sentinels. Each sentinel cage, containing 4 mice, surveys a maximum of 280 cages. Each week, each sentinel cage receives approximately 15 mL dirty bedding from 40 colony cages, alternating weekly to ensure all cages are surveyed over a 7-week period. One sentinel mouse per cage is sampled every 8 weeks, with blood collected for serology and feces/pelt swabs for PCR. Survival blood collection is performed using a sterile 25g needle from the tail vein, submandibular vein, or submental vein. One sentinel mouse per cage is euthanized at 6- and 12-months post-placement for additional testing, including blood collection for serology (cardiac puncture post-CO_2_ asphyxiation euthanasia), pelt and large intestinal content examination for ecto- and endoparasites, and gross necropsy with subsequent histology if lesions identified. Every two months for 12 months sentinels are tested for fur mites and pinworms (above), as well as *D. musculi, S. muris*, MHV, Sendai virus, MPV, MVM, PVM, TMEV, EDIM, MNV, MuAstV2, Reovirus-3, *M. pulmonis, R. pneumotropicus, C. bovis, S. aureus, Helicobacter spp*., and SFB. In addition, we test them for Ectromelia virus, LCMV, K-Virus, MAD1/2, Polyoma virus, MKPV, *C. rodentium, Salmonella* spp., *K. pneumoniae/oxytoca, S. pneumoniae*, Beta-hemolytic *Strep*., *P. aeruginosa, and Pneumocystis spp*. every six months and for MCMV, MTV, Hantaan virus, *Filobacterium rodentium, C. pilliforme, C. kutscheri, S. moniliformis, E. cuniculi, G. muris, E. muris*, and *T. muris* every 12 months. Testing for the presence of Cm was added to the 6 and 12-month routine soiled bedding colony health surveillance testing panels in December 2021.

### Fecal Collection

Fecal pellets for PCR assay or microbiome analysis were collected from the soiled bedding, directly from the animal, or both. When collected directly, animals were lifted by the base of the tail and allowed to grasp onto the wire-bar lid. Animals were gently restrained in this manner and a sterile 1.5 mL Eppendorf tube was placed underneath the anus for fecal collection directly into tube. If at the end of a 30 second period the animal did not defecate, it was returned to the cage and allowed to rest for at least 2 minutes and collection reattempted until a sample was produced. A total of 2 fecal pellets were collected for each microbiome sample, and 10 fecal pellets were collected for each PCR sample.

### *Chlamydia muridarum* PCR Assay and Amplicon Sequencing

A proprietary real-time fluorogenic 5′ nuclease PCR assay specifically targeting a 425 bp region Cm 23S rRNA was used to determine the presence of genomic DNA in samples. Samples that amplified during initial testing were retested using DNA isolated from a retained lysate sample to confirm the original finding. A positive result was reported when the retested sample was confirmed positive. To monitor for successful DNA recovery after extraction and to assess whether PCR inhibitors were present, an exogenous nucleic acid recovery control assay was added to each sample after the lysis step and prior to magnetic nucleic acid isolation. The concentration of eluted nucleic acid in mock extracted samples (no sample material) is calibrated to approximately 40 copies of exogenous DNA/uL and compared with a 100-copy system suitability control. A second real-time fluorogenic 5’ nuclease PCR assay is used to target the exogenous template, to serve as a sample suitability control and is performed simultaneously with the Cm assay. Nucleic acid recovery control assays for samples that demonstrated greater than a log10 loss of template copies compared with control wells were diluted 1:4 and retested, reextracted or both prior to accepting results as valid. A 100-copy/reaction positive control plasmid template containing the Cm target template was co-PCR amplified with the test sample to demonstrate master mix and PCR amplification equipment function. PCR products from select samples were purified and sequenced using the Sanger method. Sequence results were further processed by trimming the sequencing primers and any undetermined nucleotide bases. The clean consensus was analyzed by comparing it to available Cm sequences in GenBank.

### Pathology

Following euthanasia by CO_2_ asphyxiation, a complete necropsy was performed, gross lesions were recorded, fresh samples of lungs were collected for aerobic culture, and all tissues including heart, thymus, lungs, liver, gallbladder, kidneys, pancreas, stomach, duodenum, jejunum, ileum, cecum, colon, lymph nodes (mandibular, mesenteric), salivary glands, skin (trunk and head), urinary bladder, uterus, cervix, vagina, ovaries, oviducts, adrenal glands, spleen, thyroid gland, esophagus, trachea, spinal cord, vertebrae, sternum, femur, tibia, stifle joint, skeletal muscle, nerves, skull, nasal cavity, oral cavity, teeth, ears, eyes, pituitary gland, and brain were fixed in 10% neutral buffered formalin. After fixation, the skull, spinal column, sternum, femur and tibia were decalcified in a formic acid and formaldehyde solution (Surgipath Decalcifier I, Leica Biosystems). Tissues were then processed in ethanol and xylene and embedded in paraffin in a tissue processor (Leica ASP6025, Leica Biosystems). Paraffin blocks were sectioned at 5 μm thickness, stained with hematoxylin and eosin (H&E), and examined by a board-certified veterinary pathologist (SC). Scrolls of lung (20 μm) were obtained from paraffin blocks for bacterial genome sequencing (below).

Peripheral blood was collected from a subset of mice by cardiac puncture. For hematology, blood was collected into tubes containing EDTA. Automated analysis was performed on an automated hematology analyzer (IDEXX Procyte DX, Columbia, MO) and the following parameters were determined: white blood cell count, red blood cell count, hemoglobin concentration, hematocrit, mean corpuscular volume, mean corpuscular hemoglobin, mean corpuscular hemoglobin concentration, red blood cell distribution width standard deviation and coefficient of variance, reticulocyte relative and absolute counts, platelet count, platelet distribution width, mean platelet volume, and relative and absolute counts of neutrophils, lymphocytes, monocytes, eosinophils, and basophils. For serum chemistry, blood was collected into tubes containing a serum separator, the tubes were centrifuged, and the serum was obtained for analysis. Serum chemistry was performed on an automated analyzer (Beckman Coulter AU680, Brea, CA) and the concentration of the following analytes was determined: alkaline phosphatase, alanine aminotransferase, aspartate aminotransferase, creatine kinase, gamma-glutamyl transpeptidase, albumin, total protein, globulin, total bilirubin, blood urea nitrogen, creatinine, cholesterol, triglycerides, glucose, calcium, phosphorus, chloride, potassium, and sodium. The Na/K and the albumin/globulin ratios were calculated.

### Immunohistochemistry (IHC)

All tissues evaluated by histopathology were screened for *Chlamydia* MOMP using a technique optimized and validated by MSK’s Laboratory of Comparative Pathology. Formalin-fixed paraffin-embedded sections were stained using an automated staining platform (Leica Bond RX, Leica Biosystems). Following deparaffinization and heat-induced epitope retrieval in a citrate buffer at pH 6.0, the primary antibody against *Chlamydia* MOMP (NB100-65054, Novus Biologicals, Centennial, CO) was applied at a dilution of 1:500. A rabbit anti-goat secondary antibody (Cat. No. BA-5000, Vector Laboratories, Burlingame, CA) and a polymer detection system (DS9800, Novocastra Bond Polymer Refine Detection, Leica Biosystems) was then applied to the tissues. The 3,3’-diaminobenzidine tetrachloride (DAB) was used as the chromogen, and the sections were counterstained with hematoxylin and examined by light microscopy. Reproductive tracts from TLR3-deficient mice experimentally infected with *Chlamydia muridarum* strain Nigg were used as positive control.^5^

### In Situ Hybridization (ISH)

Select IHC-positive tissues were included for ISH analysis. The target probe was designed to detect region 581-617 of *Chlamydia muridarum* str. Nigg complete sequence, NCBI Reference Sequence NC_002620.2 (1039538-C1; Advanced Cell Diagnostics, Newark, CA). The target probe was validated on reproductive tracts from mice experimentally inoculated with *Chlamydia muridarum* strain Nigg.^5^ Slides were stained on an automated stainer (Leica Bond RX, Leica Biosystems) with RNAscope 2.5 LS Assay Reagent Kit-Red (322150, Advanced Cell Diagnostics) and Bond Polymer Refine Red Detection (DS9390, Leica Biosystems). Control probes detecting a validated positive housekeeping gene (mouse peptidylprolyl isomerase B, *Ppib* to confirm adequate RNA preservation and detection; 313918, Advanced Cell Diagnostics) and a negative control, *Bacillus subtilis* dihydrodipicolinate reductase gene (dapB to confirm absence of nonspecific staining; 312038, Advanced Cell Diagnostics) were used. Positive RNA hybridization was identified as discrete, punctate chromogenic red dots under bright field microscopy.

### *Chlamydial muridarum* 23S rRNA Amplicon Sequencing

DNA was extracted (MagMAX Total Nucleic Acid Isolation Kit, Life Technologies, Carlsbad, CA using a Qiagen robotic extraction station, Thermo Labsystems, Franklin, MA) from pulmonary tissue scrolls from an NSG mouse used as a contact sentinel according to the manufacturers’ instructions. A region of the 23S rRNA gene was amplified using HotStar Taq polymerase (Qiagen) with the primers CHLSE37 (5′ CTTGGCATTGACAGGCGA 3′) and CHLSE461 (5′ GGAGAGTGGTCTCCCCAGATT 3′), which generated an expected product of 425 bp; primer nomenclature is based on the nucleotide positions of a Cm reference sequence (GenBank accession no., U68436) and was designed to detect within the genus of Chlamydia. Thermal cycling conditions were: denaturation at 95°C for 15 min, followed by an initial 5 cycles of denaturation at 94°C for 30 s, primer annealing at 64°C for 30 s, and extension at 72°C for 60 s. The annealing temperature was then decreased to to58°C for 35 cycles; all other parameters remained as described for the initial 5 cycles. PCR products were separated on 2% agarose gels, eluted from the gel (Minelute Kit, Qiagen), and sequenced (ABI 3130XL DNA sequencer, Tufts University Core Facility, Boston, MA).

### Isolation and Growth of Chlamydiae

Samples (lung, cecum, and feces) from an NSG contact sentinel were macerated and placed in sucrose-phosphate-glutamic acid buffer (pH 7.2), frozen at -80°C, and shipped on dry ice to Midwestern University where they were again held at -80°C until isolations were conducted. Methods have been previously described.^29^ Briefly, all samples were thawed, sonicated and centrifuged (400 x g for 10 minutes at 4° C) to pellet cellular and other debris. The pelleted debris was discarded, and the supernatants were diluted in Dulbecco’s modified essential medium (DMEM, containing 584 mg/L glutamine, 4.5g/L glucose). Fecal and cecum samples were then filtered through 0.22 μM sterile filter. Samples were then inoculated on confluent monolayers of HeLa 229 in 24 well tissue culture plates at 500 μl per well. Inoculated 24 well plates were then centrifuged at 1100 g for 1 hour at 37 ° C. Following centrifugation, the plates were incubated for an additional 1 hour at 37°C in 5% CO_2_. The inocula were then aspirated and replaced with chlamydial growth media (DMEM containing, 10% fetal calf serum, cycloheximide 0.5 μg/ml) and antibiotics (vancomycin 100 µg/ml, gentamicin 50 µg/ml, and amphotericin B 1.25 µg/ml) to inhibit growth of mouse microbial flora.^29^ Monolayers were then monitored for inclusion formation by scanning twice daily by inverted microscopy. In some cases, initial dilutions were too low and chlamydial cytotoxicity was observed which we believe was attributable to chlamydial growth, as described elsewhere.^2^ In other cases, inclusions were observed by 36-, 48-, or 72-hours post-inoculation and the 24 well plate flask was fixed in cold methanol for visualization of inclusions by indirect immunofluorescence assay (IFA).^8^ Duplicate plates were run in parallel for expanding isolates and confirming chlamydia detection by 16S rRNA PCR as previously described.^41^

### Fecal Microbiome Analysis

Collection kits (Transnetyx Microbiome, Transnetyx, Cordova, TN, USA) containing barcoded sample collection tubes were used. Fecal samples (2 pellets from each mouse) were placed in separate tubes containing DNA stabilization buffer to ensure reproducibility, stability, and traceability, and shipped for DNA extraction, library preparation, and sequencing (Transnetyx). DNA extraction was performed using the Qiagen DNeasy 96 PowerSoil Pro QIAcube HT extraction kit and protocol for reproducible extraction of inhibitor-free, high molecular weight genomic DNA that captures the true microbial diversity of stool samples. After DNA extraction and quality control, genomic DNA was converted into sequencing libraries using the KAPA HyperPlus library preparation protocol optimized for minimal bias. Unique dual indexed adapters were used to ensure that reads and/or organisms were not mis-assigned. Thereafter, the libraries were sequenced using the Illumina NovaSeq instrument and protocol via the shotgun sequencing method (a depth of 2 million 2×150 bp read pairs), which enables species and strain level taxonomic resolution. Raw data (in the form of FASTQ files) was analyzed using the One Codex analysis software and analyzed against the One Codex database consisting of >115K whole microbial reference genomes. The One Codex Database consists of ∼114K complete microbial genomes, including 62K distinct bacterial genomes, 48K viral genomes, and thousands of archaeal and eukaryotic genomes. Human and mouse genomes are included to screen out host reads. The database is assembled from both of public and private sources, with a combination of automated and manual curation steps to remove low quality or mislabeled records. Every individual sequence (NGS read or contig) is compared against the One Codex database by exact alignment using k-mers where k=31. Based on the relative frequency of unique k-mers in the sample, sequencing artifacts are filtered out of the sample to eliminate false positive results caused by contamination or sequencing artifacts. The relative abundance of each microbial species is estimated based on the depth and coverage of sequencing across every available reference genome.

## Results

### Index Cases

Laboratory A: The initial index cases were 2 experimentally naïve, 1.5-year-old, female, C57Bl/6-Foxp3^tm1Flv^/J; C57Bl/6/Foxp3^CreER^R26^tdTomato^ mice that were noted to have unexpected inflammatory infiltrates on histology. Microscopically, 1 of the 2 mice (C57Bl/6/Foxp3^CreER^R26^tdTomato^) examined had perivascular and peribronchiolar lymphoplasmacytic aggregates (Figure 1A and B). Pulmonary FFPE scrolls from the affected lung assayed by real time PCR targeting of the chlamydial 23S rRNA gene followed by sequencing of the resulting amplicon yielded chlamydial DNA with 98.22% identity with Cm isolate CM001 (GenBank accession: CP027217.1). Subsequently, bronchial epithelium staining for *Chlamydia* MOMP by IHC was detected in an affected airway (Figure 1C). Additional spontaneous and/or incidental age-related lesions found in these mice included: mild, lymphoplasmacytic enteritis (n=2); mild cecal tritrichomoniasis (n=2); mild multifocal hepatocellular necrosis associated with neutrophilic and histiocytic infiltrates (n=2); mild, lymphoplasmacytic colitis (n=1), lymphocytic interstitial nephritis (n=1); lymphocytic aggregates in the mesentery, mediastinum and the salivary glands (n=1); subcapsular spindle cell hyperplasia in the adrenal gland (n=1). There were no microscopic lesions noted in the cecum and colon of these mice (Figure 1D); however, chlamydial antigen was detected in moderate amounts within surface cecal and colonic epithelial cells associated with gut associated lymphoid tissue (Figure 1E). Chlamydial antigen was not detected in other organs assayed (heart, thymus, kidneys, adrenal glands, liver, gallbladder, spleen, pancreas, stomach, duodenum, jejunum, ileum, salivary glands, lymph nodes, trachea, esophagus, reproductive tract, mammary glands, skin, bones, skeletal muscle, femoral and spinal nerves, teeth, pituitary gland, and brain). Chlamydial 23S rRNA was also detected in select cecal and colonic epithelial cells by ISH (Figure 1F). Red and white blood cell counts and serum chemistry results from these mice were unremarkable.

**Figure 1.**
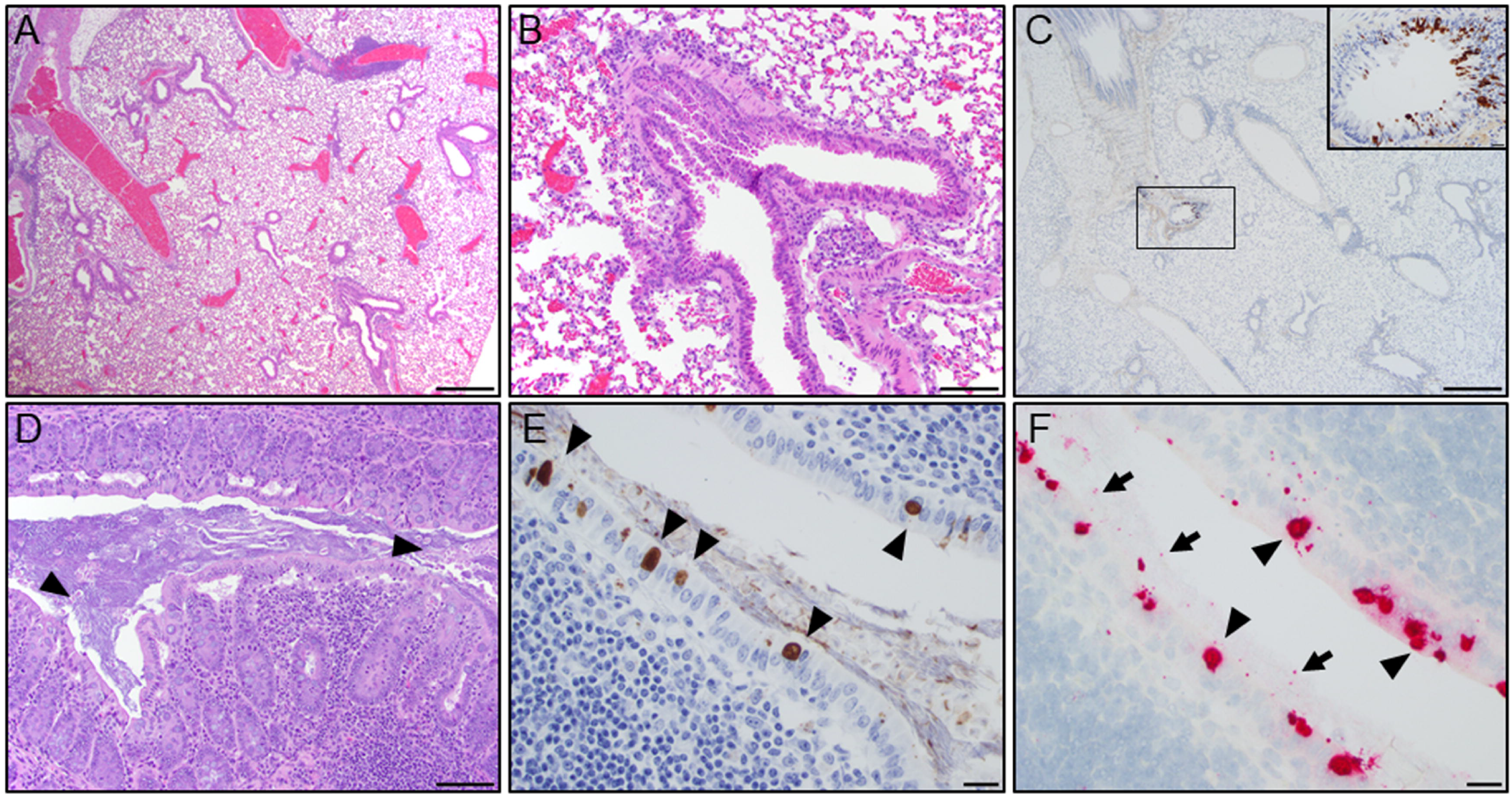
Representative histopathology of the lung and cecum from a 1.5-year-old, female, C57Bl/6/Foxp3^CreER^R26^tdTomato^ mouse. **A**. Multifocally, peribronchiolar and perivascular spaces are infiltrated by small to moderate clusters of lymphocytes and histiocytes (scale bar = 500µm). **B**. High magnification field shows lymphocytic and histiocytic infiltrates around a bronchiole (scale bar = 100µm). **C**. IHC of the lung demonstrating focal detection of chlamydial MOMP antigen in bronchiolar epithelial cells (scale bars = 100µm; 20 µm - brown staining in inset). **D**. Representative section of normal mucosa and gut-associated lymphoid tissue (GALT) in the cecum with low numbers of luminal *T. muris* (arrowheads and inset; scale bars = 100µm). **E**. IHC of the cecum demonstrating detection of intracytoplasmic chlamydial inclusions in surface epithelial cells (arrowhead; scale bar = 20 µm). **F**. ISH demonstrates positive staining (red) for Cm mRNA within the cecal epithelium (arrowhead) and lumen (arrow; scale bar = 20 µm).

Laboratory B: The second set of index cases were 2, 4-week-old, female Cd23-Cre Gen1^-/-^ Mus81^fl/fl^ mice that had presented for alopecia and runting. Microscopically, skin lesions were consistent with a spontaneous follicular dystrophy known to occur in mice of this genetic background.^39^ These mice also demonstrated a mild, multifocal, chronic lymphoplasmacytic nasopharyngitis, tracheitis, bronchiolitis and bronchitis. IHC revealed focal staining of chlamydial MOMP in bronchiolar epithelial cells in normal bronchioles and bronchioles surrounded by lymphoplasmacytic infiltrates in both mice as well in surface colonic epithelium in 1 of 2 mice. Pulmonary FFPE scrolls from 1 of 2 mice were submitted for genus-specific PCR and amplicon sequencing (as described for mice from Laboratory A) and yielded similar results. Other microscopic findings noted in these mice included neutrophilic otitis media (n=2), small intestinal spironucleosis (n=2), subacute nasopharyngitis (n=2) and minimal to mild tracheitis and/or hepatitis (n=1).

### Analysis of Imported Mice for *Chlamydia muridarum*

Approximately 33% of the 58 groups of mice imported from academic institutions, pharmaceutical/biotechnology companies, and non-approved commercial breeders assessed for Cm infection by fecal PCR were positive (Table 1). The imported group was classified “positive” when Cm was detected via PCR on samples collected during both QA and QB. There were several groups (n=4) in which only samples collected during QB were positive. In 3 of 4 of these cases, the laboratory had introduced Cm positive animals from their colony for breeding during quarantine and therefore these groups were classified as “negative.” The remaining case was unrelated to the introduction of colony animals; a subsequent additional (3^rd^) positive PCR assay was conducted, and the group was classified as “positive.” The initial negative result was presumed to be a false negative. The 19 positive groups were imported from 16 different institutions. While the preponderance of the groups received were imported from other US institutions, a group of mice imported from Germany was also positive. No mice imported during this period were observed to have any clinical abnormalities.

**Table 1:** Detection of *Chlamydia muridarum* in imported mice

|  | Sample Totals |  |  | Institution Totals |  |  |
| --- | --- | --- | --- | --- | --- | --- |
|  | # Samples | # Positive | % Positive | # Institutions | # Positive | % Positive |
| Europe | 8 | 1 | 12.5% | 7 | 1 | 14.3% |
| Austria | 1 | 0 | 0.0% | 1 | 0 | 0.0% |
| Denmark | 1 | 0 | 0.0% | 1 | 0 | 0.0% |
| France | 1 | 0 | 0.0% | 1 | 0 | 0.0% |
| Germany | 3 | 1 | 33.3% | 2 | 1 | 50.0% |
| UK | 2 | 0 | 0.0% | 2 | 0 | 0.0% |
| Asia | 2 | 0 | 0.0% | 2 | 0 | 0.0% |
| Japan | 2 | 0 | 0.0% | 2 | 0 | 0.0% |
| Northeast US | 28 | 10 | 35.7% | 15, 1<sup>α</sup> | 9, 0<sup>α</sup> | 60.0%, 0.0%<sup>α</sup> |
| Connecticut | 1 | 1 | 100.0% | 1 | 1 | 100.0% |
| Distr. Columbia | 1 | 0 | 0.0% | 1 | 9 | 0.0% |
| Maryland | 5 | 1 | 20.0% | 4 | 1 | 25.0% |
| New York | 16 | 5 | 31.3% | 7, 1<sup>α</sup> | 5, 0<sup>α</sup> | 71.4%, 0.0%<sup>α</sup> |
| Pennsylvania | 5 | 3 | 60.0% | 3 | 2 | 66.7% |
| Midwest US | 6 | 3 | 50.0% | 3 | 2 | 66.7% |
| Missouri | 5 | 2 | 40.0% | 2 | 1 | 50.0% |
| Wisconsin | 1 | 1 | 100.0% | 1 | 1 | 100.0% |
| South US | 8 | 2 | 25.0% | 7 | 2 | 28.6% |
| Alabama | 1 | 0 | 0.0% | 1 | 0 | 0.0% |
| North Carolina | 3 | 0 | 0.0% | 2 | 0 | 0.0% |
| Tennessee | 1 | 0 | 0.0% | 1 | 0 | 0.0% |
| Texas | 3 | 2 | 66.7% | 3 | 2 | 66.7% |
| West US | 6 | 3 | 50.0% | 3, 1<sup>β</sup> | 2, 0<sup>β</sup> | 66.7%, 0.0%<sup>β</sup> |
| California | 6 | 3 | 50.0% | 3, 1<sup>β</sup> | 2, 0<sup>β</sup> | 66.7, 0.0%<sup>β</sup> |
| **Total** | 58 | 19 | 32.8% | 37, 1<sup>α</sup>, 1<sup>β</sup> | 16, 0<sup>α</sup>, 0<sup>β</sup> | 43.2%, 0.0%<sup>α</sup>, 0.0%<sup>β</sup> |

Results are from academic colonies unless otherwise indicated. α = non-commercial breeder; β = biotechnology company

### Colony Surveillance (Sentinels) for *Chlamydia muridarum*

Of the 97 animal holding rooms from 3 institutions that were surveyed by examining the associated soiled bedding sentinel mice for Cm via fecal PCR, 61 (62.9%) were positive representing all 8 of the 8 (100%) surveyed vivaria (Table 2). Of the 90 rooms surveyed that received imported mice following testing for excluded agents and release, 61 (67.8%) were positive for Cm. None of the 7 holding rooms that require receipt from commercial vendors or C-section rederivation of imported mice were Cm positive.

**Table 2:** Detection of *Chlamydia muridarum* in soiled bedding sentinels

| Facility | Rm. Type | # Rooms | # Positive | % Positive |
| --- | --- | --- | --- | --- |
| A |  | 18 | 13 | 72.2% |
|  | STD | 16 | 13 | 81.3% |
|  | R/V | 2 | 0 | 0.0% |
| B |  | 13 | 8 | 61.5% |
|  | STD | 13 | 8 | 61.5% |
| C |  | 3 | 3 | 100.0% |
|  | STD | 3 | 3 | 100.0% |
| D |  | 7 | 4 | 57.1% |
|  | STD | 6 | 4 | 66.7% |
|  | R/V | 1 | 0 | 0.0% |
| E |  | 36 | 19 | 52.8% |
|  | STD | 32 | 19 | 59.4% |
|  | R/V | 4 | 0 | 0.0% |
| F |  | 12 | 8 | 66.7% |
|  | STD | 12 | 8 | 66.7% |
| G |  | 5 | 3 | 60.0% |
|  | STD | 5 | 3 | 60.0% |
| H |  | 3 | 3 | 100.0% |
|  | STD | 3 | 3 | 100.0% |
| **Total** |  | 97 | 61 | 62.9% |

STD = Standard sources; animal holding rooms containing mice obtained from commercial breeders and non-commercial sources (released after testing for specific agents). R/V = Rederived/Vendor: animal holding rooms receiving rederived mice from non-commercial sources (redерived into axenic state and reassociated with defined, vendor-specific flora) and/or commercial breeders.

### Exposure of NSG Mice to *Chlamydia muridarum* Shedding Imported Mice

One of the 2 NSG mice that was cohoused with Cm shedding (confirmed via PCR) imported mice and 4 of the 8 NSG mice that were cohoused with these NSG contacts that continued to receive soiled bedding from the imported mice became acutely ill 3 weeks after the initial placement of the 2 NSG contact sentinels and were euthanized. The mice were lethargic, hunched, had unkempt hair coats and had lost weight. The remaining 5 NSG mice (including the second contact sentinel and 4 remaining soiled bedding samples) were euthanized 1 week later when similar clinical signs developed. The initial 2 contact sentinels had a moderate to severe, multifocal to coalescing, chronic, histiocytic and neutrophilic bronchointerstitial pneumonia. Bronchiolar lumens contained degenerate neutrophils, necrotic cells, karyorrhectic debris, fibrin, and proteinaceous eosinophilic fluid (Figure 2A). Bronchiolar epithelial cells often contained clear intracytoplasmic vacuoles with pale-basophilic inclusions compatible with chlamydial inclusions (Figure 2A). Intracytoplasmic inclusions were also noted in macrophages and alveolar epithelial cells within regions of peribronchiolar and alveolar inflammation. Peribronchiolar and alveolar spaces were infiltrated by large numbers of macrophages and neutrophils associated with edema or contained pyknotic cells, karyorrhectic debris and fibrin. Additional microscopic findings in these mice included a minimal to mild, multifocal, subacute neutrophilic and histiocytic typhlocolitis with crypt and goblet cell hyperplasia and crypt dilatation. Chlamydial MOMP antigen was detected by IHC in many surface epithelial cells in histologically normal regions of the small and large intestine (Figure 2D and 2F) as well as in cecal segments with the aforementioned inflammation and in histologically normal segments of the trachea, nasopharynx, and lungs of both mice. A mouse demonstrated positive staining in the nasal cavity, and another in the spleen. Chlamydial mRNA was also detected in the cecal and colonic epithelium (Figure 2E), as well as in tracheal and lung epithelial cells of both mice by ISH. There was good correlation between the inclusions noted in the lung on H&E with IHC and ISH staining, detecting chlamydial MOMP antigen and mRNA, respectively, in affected bronchiolar epithelial cells and regions of peribronchiolar and alveolar inflammation (Figure 2B and C). Chlamydia MOMP antigen and mRNA were also detected in the affected bronchiolar lumens admixed with degenerate neutrophils, necrotic debris, and proteinaceous material as described above. Pulmonary FFPE scrolls from the 2 mice were Cm PCR positive. Aerobic pulmonary cultures from both mice were negative. A moderate leukocytosis (5.50-934 ×10^3^/µL, reference range 0.94-4.68 ×10^3^/µL) characterized by a moderate to marked neutrophilia (5.18-8.77 ×10^3^/µL, reference range 0.54-3.16 ×10^3^/µL) was also observed in these mice.

**Figure 2.**
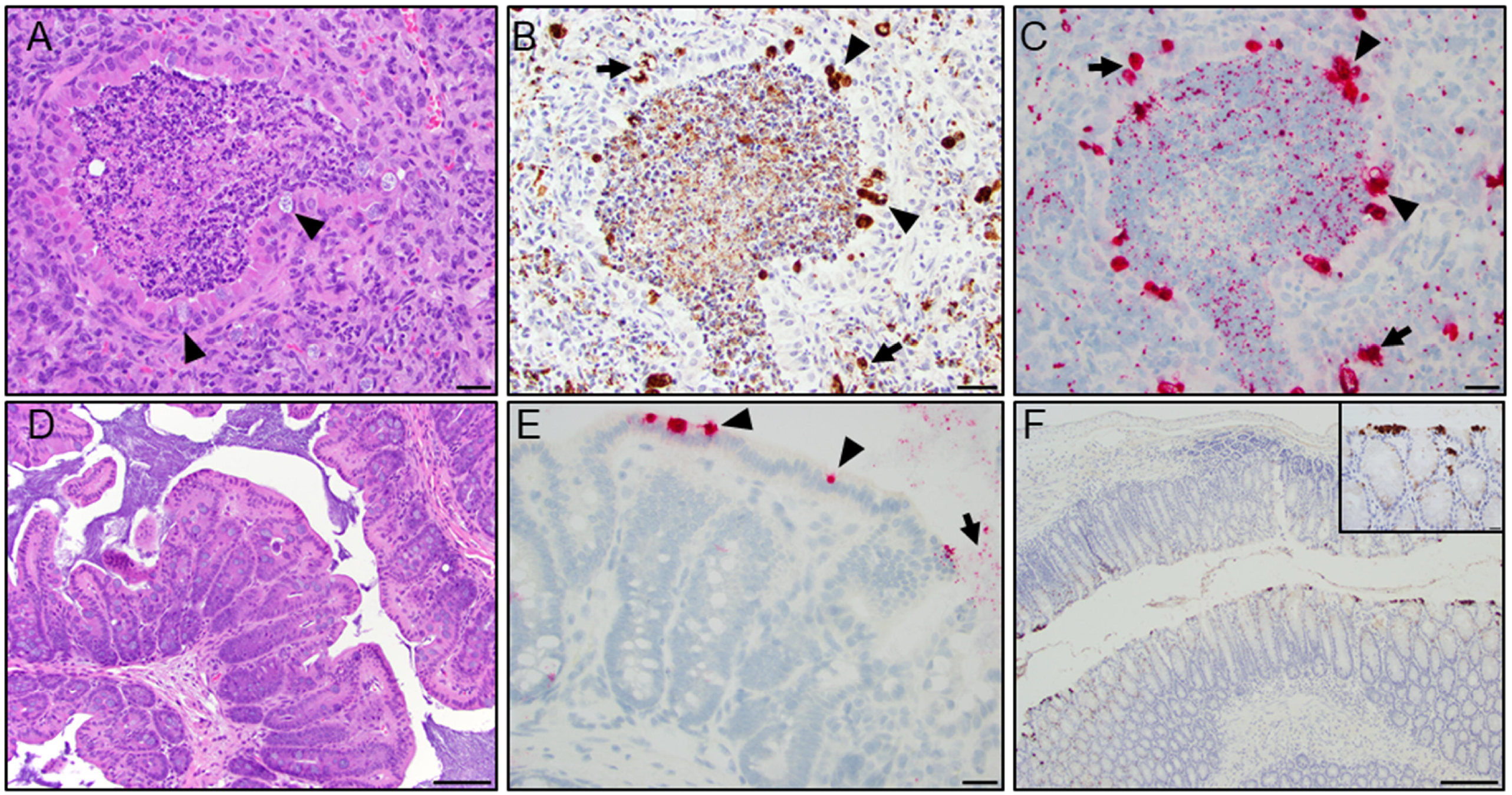
Histopathology of the lung and large intestine from a 1-year-old, female, NSG mouse. **A**. Representative airway demonstrating a bronchiolar and alveolar inflammation characterized by luminal neutrophilic infiltration mixed with necrotic debris and proteinaceous material. Multifocally, bronchiolar epithelial cells exhibit intracytoplasmic clear vacuoles with pale-basophilic structures compatible with Chlamydial inclusions (arrowhead and inset). Peribronchiolar and alveolar space is infiltrated with moderate numbers of macrophages and neutrophils intermixed with reactive fibroblasts (scale bar = 50 µm). **B**. IHC of the lung demonstrating detection of chlamydial MOMP antigen in bronchiolar epithelial cells (arrowhead; brown staining) and areas with peribronchiolar inflammation (arrow, scale bar = 50 µm). **C**. ISH demonstrates positive staining (red) for Cm mRNA in bronchiolar epithelial cells (arrowhead) and areas of peribronchiolar inflammation (arrow; scale bar = 50 µm). **D**. Representative H&E-stained section of a normal cecal wall (scale bars = 100µm). **E**. High magnification field demonstrates ISH signal (red staining) in the cecal epithelium (arrow) and lumen (arrowhead; scale bar = 50µm). **F**. IHC of descending colon demonstrating detection of intracytoplasmic chlamydial MOMP antigen in surface epithelial cells (inset - brown staining; scale bars = 20 -200µm).

### *Chlamydial muridarum* 23S rRNA Amplicon Sequencing

Sequence analysis of the amplified DNA 387 bp fragment from a segment of the 23S rRNA gene showed 100% identity with 22 Cm Nigg strain isolates (GenBank accessions: CP027217.1, CP027216.1, CP027215.1, CP027214.1, CP027213.1, CP027212.1, CP027211.1, CP027210.1, CP027209.1, CP027208.1, CP027207.1, CP027206.1, CP007217.1, CP009760.1, CP009609.1, CP009608.1, CP007276.1, CP006975.1, CP006974.1, NR_076163.1, CP063055.1, and AE002160.2) as well as a single Cm isolate listed only as MoPn (GenBank accession: U68436.4) and a Cm isolate listed as SFPD, noted as isolated in a hamster (GenBank accession: U68437.2). Notably, there was no sequence homology with the single strain Weiss whole genome sequence (GenBank accession: NZ_ACOW00000000.1); however, close examination revealed the sequence was incomplete and lacked the entire amplified DNA 387 bp fragment.

### Assessment of Commercial Breeders, Pet Store and Wild Mice for *Chlamydia muridarum*

*Chlamydia muridarum* was not detected in feces collected from any of the mice received directly from any of the 55 production housing rooms, located in 5 US states and 2 Canadian provinces surveyed from 4 commercial breeders. All 3 samples from Pet Shop A, collected at 2 distinct time points, were positive for Cm. Samples from pet shops B and C were negative. The fecal sample from the single wild mouse population was negative.

### Isolation of *Chlamydia muridarum*

*Chlamydia muridarum* was isolated from the lung, cecum and feces of an NSG mouse that was cohoused with imported mice shedding Cm. Cytotoxicity was observed in HeLa 229 cell cultures inoculated with 1:10 dilutions of filtered supernatants from lung, cecum and feces by 24 hours post-inoculation. Cytotoxicity is a well described observation for chlamydial species that carry a cytotoxin orthologous to the clostridial cytotoxin of which Cm is known to carry 3 copies.^2^ Classic chlamydial inclusions were observed 24 hours post-inoculation when higher dilutions of samples were cultured on HeLa 229 cell monolayers. These inclusions often displayed the typical Brownian movement seen with Cm infection and stained by IFA (Figure 3). Cm was also confirmed in these infected monolayers by PCR for Cm 16S rRNA.

**Figure 3.**
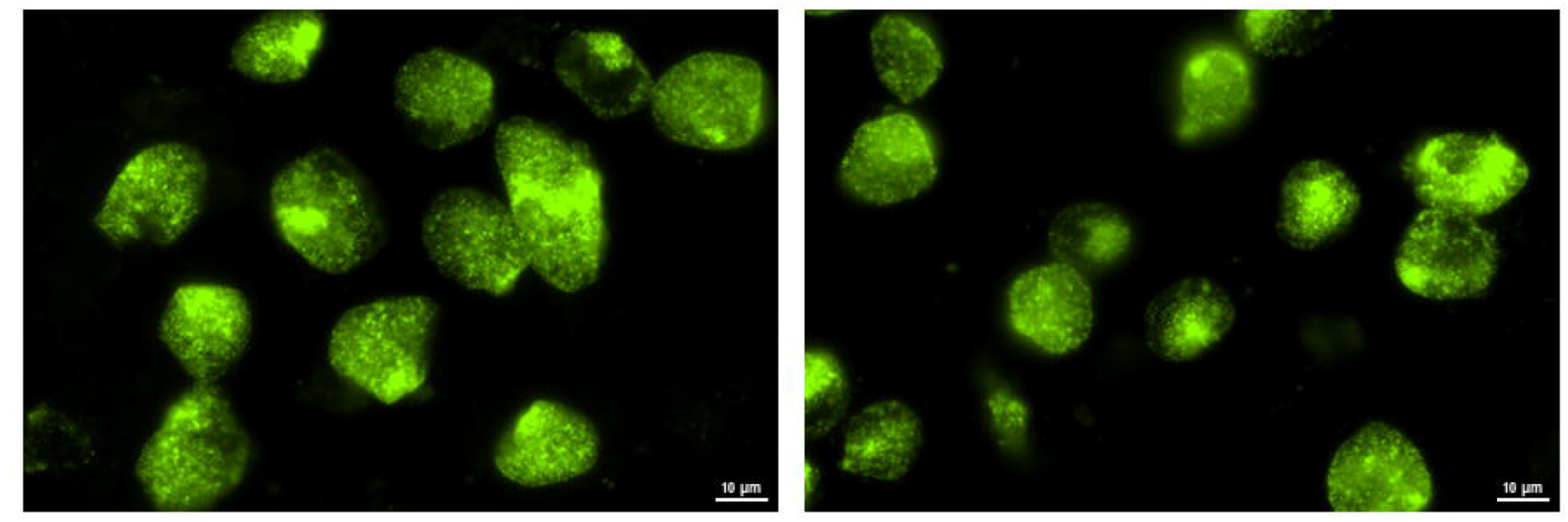
Fluorescent images of *Chlamydia*-infected cells. HeLa 229 cells were infected with *Chlamydia* spp. isolated from an NSG mouse cecum sample. At 30 hours post-infection, cells were fixed and stained with FITC-labeled anti-*Chlamydia* spp. antibody (green) and analyzed using an EVOS™ FL Auto Imaging fluorescent microscope (Life Technologies).

### Prevalence of *Chlamydia muridarum* in Diagnostic Samples

The prevalence of Cm in diagnostic samples from various institutions and source materials was 16.2% (146 of 900 samples tested), with 9.9% (54 of 546) of the samples collected directly from mice, 29.7% (85 of 286) of the environmental samples, and 12.1% (7 of 58) of combination animal/environmental samples testing positive (Table 3). None of the biologics nor the samples collected from rodent production facilities tested positive. Samples from academic research facilities (133 of 519, 25.6%) were much more likely to test positive as compared to samples from pharmaceutical (5/249, 2.0%) or contract research facilities (8/76, 10.5%).

**Table 3:** Prevalence of *Chlamydia muridarum* in diagnostic samples

| Sample Type | # Samples | # Positive | % Positive |
| --- | --- | --- | --- |
| Animal | 546 | 54 | 9.9% |
| Academic Research | 282 | 46 | 16.3% |
| Pharmaceutical | 162 | 0 | 0.0% |
| Contract Research | 71 | 8 | 11.3% |
| Rodent production | 31 | 0 | 0.0% |
| Environmental | 286 | 85 | 29.7% |
| Academic Research | 207 | 80 | 38.7% |
| Pharmaceutical | 72 | 5 | 6.9% |
| Contract Research | 0 | 0 | 0.0% |
| Rodent production | 7 | 0 | 0.0% |
| Anim./Enviro. Combination | 58 | 7 | 12.1% |
| Academic Research | 27 | 7 | 25.9% |
| Pharmaceutical | 8 | 0 | 0.0% |
| Contract Research | 5 | 0 | 0.0% |
| Rodent production | 18 | 0 | 0.0% |
| Biologics | 10 | 0 | 0.0% |
| Academic Research | 3 | 0 | 0.0% |
| Pharmaceutical | 7 | 0 | 0.0% |
| Contract Research | 0 | 0 | 0.0% |
| Rodent production | 0 | 0 | 0.0% |
| **Total** | **900** | **146** | **16.2%** |

### Fecal Microbiome analysis

The metagenomic microbiome sequencing data results for Cm are provided in Table 4. Overall, institutional prevalence of Cm abundance was 14.17% (17 of 120 institutions). An additional institution had abundance of Chlamydia at the genus, but not the species level. While the preponderance of the samples were from academic institutions (112 of 120; 93%) located in the United States (91 of 120; 76%), Cm was also detected as a component of the fecal microbiome in European colonies (3 of 23; 13%). None of the 8 commercial institutions, all of which were from North America, had Cm as a component of their intestinal flora. There were considerable differences in the number of samples assayed from institutions with Cm abundance, ranging from as few as 3 to as many as 4756 samples per institution. The sample prevalence among these institutions ranged from as low as 1% to as high as 100% testing positive. Academic institutions from which positive samples were obtained were in all regions of the US with an overrepresentation of samples submitted from the Northeast and the South.

**Table 4:** Metagenomic fecal microbiome sequence data for *Chlamydia muridarum*

| Category / Region | Sample Totals |  |  | Institution Totals |  |  |
| --- | --- | --- | --- | --- | --- | --- |
|  | # Samples | # Positive | % Positive | # Institutions | # Positive | % Positive |
| Academic | 10,903 | 383 | 3.5% | 112 | 17 | 15.0% |
| Australia | 115 | 0 | 0.0% | 5 | 0 | 0.0% |
| Europe | 728 | 47 | 6.5% | 23 | 3* | 13.0% |
| Institution A* | 8 | 2 | 25.0% |  |  |  |
| Institution B* | 39 | 7 | 18.0% |  |  |  |
| Institution C* | 52 | 38 | 73.1% |  |  |  |
| Northeast US | 976 | 223 | 22.8% | 27 | 7* | 26.0% |
| Institution D* | 10 | 8 | 80.0% |  |  |  |
| Institution E* | 11 | 11 | 100.0% |  |  |  |
| Institution F* | 36 | 6 | 16.7% |  |  |  |
| Institution G* | 68 | 32 | 47.1% |  |  |  |
| Institution H* | 524 | 166 | 31.7% |  |  |  |
| Midwest US | 1,603 | 15 | 0.9% | 20 | 1* | 5.0% |
| Institution I* | 45 | 15 | 33.3% |  |  |  |
| Southern US | 7,003 | 46 | 0.7% | 18 | 5* | 28.0% |
| Institution J* | 30 | 3 | 10.0% |  |  |  |
| Institution K* | 16 | 6 | 37.5% |  |  |  |
| Institution L* | 40 | 5 | 12.5% |  |  |  |
| Institution M* | 53 | 4 | 7.6% |  |  |  |
| Institution N* | 4,756 | 28 | 0.6% |  |  |  |
| Western US | 681 | 7 | 1.0% | 19 | 3* | 16.0% |
| Institution O* | 3 | 1 | 33.3% |  |  |  |
| Institution P* | 5 | 5 | 100.0% |  |  |  |
| Institution Q* | 82 | 1 | 1.2% |  |  |  |
| Commercial | 429 | 0 | 0.0% | 8 | 0 | 0.0% |
| Canada | 20 | 0 | 0.0% | 1 | 0 | 0.0% |
| Northeast US | 168 | 0 | 0.0% | 4 | 0 | 0.0% |
| Midwest US | 168 | 0 | 0.0% | 1 | 0 | 0.0% |
| Southern US | 73 | 0 | 0.0% | 2 | 0 | 0.0% |
| **Total** | **11,387** | **383** | **3.4%** | **120** | **17** | **14.2%** |

\*Sample details for institutions yielding positive results in each region.

## Discussion

We provide compelling evidence that Cm is moderately prevalent and globally distributed in laboratory mouse colonies in numerous academic biomedical research institutions. This is evidenced by PCR and fecal metagenomic data from 2 large sample sets, testing of imported mice from other academic colonies, and the prevalence within our institutions (MSK, Weill Cornell Medicine, and the Hospital for Special Surgery). The initial identification of Cm followed the pursuit of the etiology of pulmonary lesions, often reported by comparative pathologists as incidental, in 2 immunocompetent GEM. This was a stunning finding considering Cm had last been identified in laboratory mice in the 1940’s and has not been considered an agent of concern since. As Cm has not been considered an ‘excluded agent’, institutions, whether they be commercial breeders or end users, were not testing for its presence. The noted exception were laboratories using the bacterium in mice as a model for *Chlamydia trachomatis* infections. If present in mice obtained from commercial breeders, they would likely have detected its presence in uninfected study controls. Importantly and consistent with this finding, Cm does not appear to be a component of the gastrointestinal microbiota of commercially bred, globally distributed mice. As such, its moderate prevalence is likely unrelated to distribution of Cm infected mice by a major commercial breeder(s). This is fortunate as the prevalence of infection would likely have been considerably higher had this not been the case. The absence of Cm in commercially reared mice is not surprising, as mice bred by commercial breeders have been rederived, or are descendants of mice that have been rederived by C-section or embryo transfer and subsequently fostered onto or transferred into mice derived using a similar method.

This begs the question as to the source(s) of Cm and how has it become endemic in so many colonies across the globe. Cm could have been introduced by Cm-infected feral mice gaining access to laboratory mouse colonies. While there has been no evidence that laboratory mice were spontaneously/naturally infected since its’ last reported isolation from laboratory mice in the 1940s, there is evidence Cm is found in wild populations of other rodents, e.g., *Peromyscus* spp.^30^ Additionally, we repeatedly detected Cm in mice obtained from a pet store. These mice are likely sourced from an enzootically infected colony(ies). Interestingly, we did not find Cm in the single wild mouse population we surveyed. Evaluation of additional wild mouse populations is warranted to determine if feral mice pose a high-level risk of exposure to this bacterium. As many research facilities are infested with escaped laboratory mice, some of which likely bred with feral populations, these mice could be Cm-carriers and pose a risk to laboratory colonies. Escape of a passaged experimental isolate could also be the source. Mice experimentally infected with Cm have and continue to be used by numerous scientists as a model to study human genital chlamydial infections caused by *C. trachomatis*. Historically, at some institutions, mice experimentally infected with Cm were handled using standard husbandry practices. Therefore, there is the potential that contamination of surfaces and equipment could have led to cross contamination of institutional mouse colonies.

Regardless of how, and how frequently, Cm was introduced undetected into laboratory mouse colonies, the interinstitutional global distribution of unique GEM strains greatly exacerbated and may be the major contributor to the observed prevalence of Cm infection. This is supported by the considerable percentage of Cm positive mice imported into our institution. While some institutions rederive mouse strains imported from other institutions, the great majority of academic institutions have and continue to quarantine and test mice for excluded agents, releasing mice negative for these agents into existing colonies. We speculate that institutions using the former method are much less likely to have introduced Cm into their colonies. Those that test imported mice for specific excluded agents, an exclusion list which previously did not include Cm, are considerably more likely to have introduced Cm into their colonies. This suspicion is supported by the finding that, at our institutions, colonies which received mice that were C-section rederived on importation and/or were only from commercial vendors were Cm free. In contrast, colonies which received mice from commercial vendors and from other institutions using a test (for excluded agents) and release system were generally Cm infected. This latter scenario resembles that which occurred with *Demodex musculi*, which has also been widely distributed in association with the exchange of GEM strains.^23^

There is prior evidence of Cm’s presence in European laboratory mouse colonies. In a recent study investigating next-generation sequencing of the microbiome as a colony health surveillance tool, Cm was detected in fecal pellets from mice housed in conventional facilities.^34^ Cm was included in a list of pathogenic bacteria, however the significance of its detection was not further explored although it was described as a non-enteric pathogen.^34^ Another recent study investigating resistance to *Plasmodium yoelii* assessed the cecal microbiome of C57BL/6 mice and detected bacteria in the Chlamydia phylum, however there was no further discussion of this finding.^37^ The significance of the bacterium’s presence was apparently not recognized in these studies, which is likely a result of the dearth of published data about spontaneous Cm infection since the 1940’s.

Once introduced, it remains unclear how readily Cm is transmitted between cages of mice. Studies have previously shown that Cm can be passed from mouse-to-mouse housed in the same cage, likely by the oral route.^8,27^ While no studies have definitively demonstrated aerosol transmission of Cm, other Chlamydiae including *C. pneumoniae* are transmitted via aerosols.^31,33^ Given the conservation observed among *Chlamydia* spp., along with the historically and currently documented respiratory pathology, aerosol transmission cannot be excluded as a potential means of transmission.^6,31,35^ Chlamydial EBs are relatively hardy, having adapted for transient extracellular survival until internalization into cells can occur.^13^ This stability is further supported by its cecal and large intestinal tropism, as this indicates resistance to gastric acidity.^42^ The relative ease of distribution of Cm through soiled bedding, as reflected by the large number of soiled bedding sentinels that were fecal PCR positive at our institutions, suggests that cross-contamination is certainly plausible and probable within a vivarium. Noteworthy is also the value of soiled bedding sentinels, and presumably other methods, e.g., PCR testing of pooled soiled bedding or exhaust air duct filters, for determining whether Cm is present within a laboratory mouse colony.

Although Cm has been used extensively to model *C. trachomatis*, limited detail is known about the biology of the bacterium following natural infection of its’ presumed natural host, *Mus musculus*. Experimental infection with Cm typically involves inoculation of either the murine urogenital tract, upper respiratory tract, or on occasion, the gastrointestinal tract. Urogenital inoculation is, by far, the most frequent method of inoculation described in the literature, as Cm is used to model *C. trachomatis*-induced urogenital disease in humans. As such, its experimental use fails to model natural/spontaneous Cm infection in several ways. First, the presumed site of natural infection and colonization is the gastrointestinal tract, similar to chlamydiae in other animals.^31,42^ Secondly, urogenital inoculation often follows administration of progesterone. Finally, the infectious doses utilized in these studies are typically very large (often ranging between 10^5^ and 10^7^ inclusion forming units [IFUs]), which likely exceeds exposures that occur in natural infections. In a recent study, Yeruva, et al. orally inoculated groups of C57BL/6 mice with Cm doses ranging from 10^2^ to 10^6^ IFU, demonstrating an ID_50_ of <10^2^ IFU.^42^ Utilizing inocula that greatly exceed those to which mice may be exposed in an enzootically infected colony may not accurately model natural infection. Previous studies do shed some light on the biology of Cm infection. For instance, while Cm was originally isolated from the respiratory tract, its primary site of infection and continued replication appears to be the gastrointestinal tract as supported by experimental observations.^11,16,25,31,42^ Although Cm can acutely infect lung and urogenital tissue in mice, it establishes chronic persistent infections in the gastrointestinal tract when inoculated orally.^42^ It is quite common for chlamydia to be carried in the gastrointestinal tract in other species, an observation long known to chlamydiologists and veterinarians. Chlamydiae reside in the gastrointestinal tract for long periods of time in the absence of clinical disease in virtually all hosts, including birds and mammals. Our current observations are consistent as we detected Cm in the intestinal tract epithelium and feces of both immunocompetent and immunocompromised mice.

Experimental studies have also demonstrated that Cm infection of immunocompetent mouse strains is generally subclinical while immunodeficient strains often succumb to infection.^19,21,42^ While immunocompetent mice were historically considered to resolve infection, recent studies suggest a chronic, subclinical infection develops in which immunocompetent animals without clinical signs and normal leukograms remain culture positive and/or the presence of Cm is observed microscopically 50-265 days post-infection.^13,32,42^ Differences exist amongst immunocompetent strains with BALB/c mice sustaining higher Cm burdens, longer infections and greater respiratory pathology as compared to C57BL/6 mice.^42^ Studies indicate a Th1 immune response is principally required to control Cm infection.^4,19,26,31,42^ Furthermore, the immune response may be significantly more intense on reinfection.^31^ Immunocompromised strains, including athymic nude, SCID, and other immunodeficient transgenic strains (including IFNγ, TLR, STAT1, MHCII/CD4, and Rag1 knockouts) have been shown experimentally to develop more significant clinical signs, dissemination of Cm, and protracted courses of infection.^5,8,9,14,21,28,31^ Long-term immunological and other biological effects are likely following Cm infection, however additional work is needed to more thoroughly elucidate these changes as they may confound studies in which Cm-infected mice are utilized.

As we observed in NSG mice used as sentinels and from which Cm was isolated, and similar to the immunodeficient strains indicated above, mice with a compromised immune system appear unable to control infection with many developing pneumonia by 30 days following inoculation. ^8,17,21,28,31^ We observed significant clinical disease in severely immunocompromised NSG mice approximately 3 weeks after cohousing with Cm shedding imported mice. The clinical signs observed were described in the experimentally infected immunodeficient strains noted above. Leukogram changes were consistent with inflammation secondary to infectious processes. In the NSGs, significant pathology was observed in pulmonary sections in association with characteristic chlamydial inclusions in bronchial epithelium. Inclusions were confirmed to be *Chlamydia* spp. by IHC MOMP staining and Cm by ISH. Pulmonary lesions noted in these NSG mice share similarities with lesions reported in the lungs from TLR2 deficient mice challenged with Cm intranasally. TLR2 deficient mice exhibited a proinflammatory cytokine profile and an exaggerated neutrophilic infiltration in the airways.^17^ There was no colocalization of Cm inclusions with the inflammatory cell infiltrates of the lamina propria of the large intestine. Further, Cm was observed in histologically normal sections of the small and large intestine as well as in unaffected regions of the cecum by both IHC and ISH. Thus, it seems unlikely that these lesions were triggered by the presence of Cm. These lesions could be a result of enhanced antigenic stimulation from alterations in the mucosal epithelial lining or enhanced trafficking of leukocytes in the lamina propria. While colonization and shedding may be prolonged, especially in gastrointestinal tissues and feces, overt inflammation of the gastrointestinal tract is not associated with chronic experimental infection, and no gastrointestinal pathology was documented in our NSG mice.^22^ Cm was likely the sole etiology of the noted pulmonary pathology as no other bacteria were isolated by aerobic culture. Further exploration of Cm-induced disease in a highly immunocompromised mouse strain, such as the NSG, may be warranted as it could be valuable to model select characteristics of *C. trachomatis*-induced disease in humans.

It is noteworthy that experimental studies with Cm have utilized serially passaged strains whose kinetics or tissue tropism may differ from that of the “field” strains that are currently circulating in laboratory mouse colonies. Interestingly, the 2 commonly used Cm isolates (Nigg and Weiss) differ significantly from one another in both their in vitro and in vivo characteristics. For example, a recent study demonstrated that the Weiss isolate displayed greater virulence and a higher replicative capacity as compared to Nigg following both intranasal and genitourinary inoculations.^29^ There are morphologic differences noted with the Weiss isolate producing smaller IBs than Nigg.^29^ Finally, 11 mutations were noted between the sequenced Weiss genome and published Nigg sequences.^29^ These findings demonstrate that even among the well-documented and carefully studied Cm isolates there are significant differences in virulence, morphology, and genetics, highlighting the potential for field isolates to be similarly distinct.

Cm should be included in institutional biosecurity protocols and treated as an excluded agent as it has the potential to impact research and animal welfare. Adding Cm to a list of surveyed agents will empower institutions to understand colony prevalence and make further decisions about its’ eradication. These decisions may include rederivation on importation or treatment. There is limited published data on antimicrobial therapy for Cm in mice. Treatment of Chlamydial infections in humans and other species is generally effective with doxycycline and it was shown to be efficacious in eradicating Cm in a study investigating inhibition of chlamydial immunity secondary to antibiotic therapy.^38^ To date, we have successfully treated, as confirmed by repetitive negative fecal PCR assays, several small mouse colonies using commonly available doxycycline (625 PPM) impregnated feed, or drinking water dosed (1mg/mL) with the drug. However, additional effective antimicrobial agents will be needed as doxycycline would be contraindicated in studies utilizing murine Tet-on/off systems. Antibiotics would also likely alter the microbiome of treated animals which may be contraindicated in certain colonies and could further confound the studies in which treated mice are used.^3,18^

In conclusion, we have demonstrated that Cm has reemerged and is prevalent among academic mouse colonies. We believe this prevalence is at least partially attributable to global collaboration and the interinstitutional sharing of GEM strains. As Cm infections in many mouse strains are likely to result in a host of biological effects, it should be treated as an ‘excluded’ agent and institutional biosecurity protocols implemented to prevent its introduction. As eradication from many enzootically infected colonies will be warranted, additional studies are necessary to identify antimicrobials that when used will achieve this goal while minimizing undesirable effects that can also confound research. There is abundant need for future studies, including continued investigation of Cm’s natural transmission and the biological effects it has on its murine host.

## Acknowledgements

MSK Core Facilities are supported by MSK’s NCI Cancer Center Support Grant P30 CA008748. The authors would like to thank Sockie Jiao and the staff of the Laboratory of Comparative Pathology for their technical assistance with IHC and ISH, Lisa Eldred and Dr. Michael Palillo for sample collection and animal monitoring, Dr. Adam Michel for his assistance in sample collection and pathology, and Dr. William Shek for his assistance in data analysis and presentation.

## Abbreviations

Cm: *Chlamydia muridarum*
MoPn: mouse pneumonitis virus
RB: reticulate body
IB: inclusion body
EB: elementary body
MOMP: major outer membrane protein
GEM: genetically engineered mouse
FFPE: formalin-fixed paraffin embedded
IHC: immunohistochemistry
ISH: in-situ hybridization
NSG: NOD.Cg-*Prkdc*^scid^ Il2rg^tm1Wjl^/SzJ
MSK: Memorial Sloan Kettering
IFA: immunofluorescence
IFU: inclusion forming units
TLR: toll-like receptor

## References

1) Barron AL, Rank RG, Moses EB. 1984. Immune response in mice infected in the genital tract with mouse pneumonitis agent (Chlamydia trachomatis biovar). Infect Immun 44: 82–85.

2) Belland RJ, Scidmore MA, Crane DD, Hogan DM, Whitmire W, McClarty G, Caldwell HD. 2001. Chlamydia trachomatis cytotoxicity associated with complete and partial cytotoxin genes. Proc Natl Acad Sci USA 98:13984–9.

3) Boynton FD, Ericsson AC, Uchihashi M, Dunbar ML, Wilkinson JE. 2017. Doxycycline induces dysbiosis in female C57Bl/6 6NCrl mice. BMC Res Notes 10: 644.

4) Brunham RC, Rey-Ladino J. 2005. Immunology of Chlamydia infection: implications for a Chlamydia trachomatis vaccine. Nat Rev Immunol 5: 149–161.

5) Carrasco SE, Hu S, Imai DM, Kumar R, Sandusky GE, Yang XF, Derbigny WA. 2017. Toll-like receptor 3 (TLR3) promotes the resolution of Chlamydia muridarum genital tract infection in congenic C57Bl/6N mice. PLoS ONE 13: e0195165.

6) Campbell J, Huang Y, Liu Y, Schenken R, Arulanandam, Zhong G. 2014. Bioluminescence imaging of Chlamydia muridarum ascending infection in mice. PLoS ONE 9: e101634.

7) Clarke I. 2001. Evolution of Chlamydia trachomatis. Ann NY Acad Sci 1230: 11–18.

8) Cotter TW, Ramsey KH, Miranpuri GS, Poulsen CE, Byrne GI. 1997. Dissemination of Chlamydia trachomatis chronic genital tract infection in gamma interferon gene knockout mice. Infect Immun 65: 2145–2152.

9) Darville T, O’Neill JM, Andrews CW, Nagarajan UM, Stahl L, Ojcius DM. 2003. Toll-like receptor-2, but not toll-like receptor-4, is essential for development of oviduct pathology in Chlamydial genital tract infection. J Immunol 171:6187–6197.

10) DeGraves FJ, Gao D, Kaltenboeck B. 2003. High-sensitivity quantitative PCR platform. BioTechniques 34: 106–115.

11) Dochez AR, Mills KC, Mulliken B. 1937. A virus disease of Swiss mice transmissible by intranasal inoculation. Proc Soc Exp Biol Med 36: 683–686.

12) Elwell C, Mirrashidi K, Engel J. 2016. Chlamydia cell biology and pathogenesis. Nat Rev Microbiol 14: 385–400.

13) Everett KD, Bush RM, Andersen AA. 1999. Emended description of the order Chlamydiales, proposal of Parachlamydiaceae fam. Nov. and Simkaniaceae fam. nov., each containing one monotypic genus, revised taxonomy of the family Chlamydiaceae, including a new genus and five new species, and standards for the identification of organisms. Int J Syst Bacteriol 49: 415–440.

14) Fan Y, Wang S, Yang X. 1999. Chlamydia trachomatis (mouse pneumonitis strain) induces cardiovascular pathology following respiratory tract infection. Infect Immun 67: 6145–6151.

15) Gogolak, F. M. 1953. The histopathology of murine pneumonitis infection and the growth of the virus in the mouse lung. J Infect Dis 92: 254–272.

16) Gordon FB, Freeman G, Glampit JM. 1938. A pneumonia-producing filtrable agent from stock mice. Proc Soc Exp Biol Med 39: 450–453.

17) He X, Nair A, Mekasha S, Alroy J, O’Connell CM, Ingalls RR. 2011. Enhanced virulence of Chlamydia muridarum respiratory infections in the absence of TLR2 activation. PLoS One 6: e20846.

18) Hou X, Zhu L, Zhang X, Zhang L, Bao H, Tang M, Wei R, Wang R. 2019. Testosterone disruptor effect and gut microbiome perturbation in mice: early life exposure to doxycycline. Chemosphere 222: 722–731.

19) Jupelli M, Guentzel MN, Meier PA, Zhong G, Murthy AK, Arulanandam BP. 2008. Endogenous IFN-gamma production is induced and required for protective immunity against pulmonary chlamydial infection in neonatal mice. J Immunol 180: 4148–55.

20) Lipman NS, Homberger FR. 2003. Rodent quality assurance testing: use of sentinel animal systems. Lab Anim 32: 36–43.

21) Magee DM, Igietseme JU, Smith JG, Bleicker CA, Grubbs BG, Schacter J, Rank RG, Williams DG. 1993. Chlamydia trachomatis pneumonia in the severe combined immunodeficiency (SCID) mouse. Regional Immunology, 5: 305–311.

22) Morrison SG, Giebel AM, Toh EC, Spencer III HJ, Nelson DE, Morrison RP. 2018. Chlamydia muridarum genital and gastrointestinal infection tropism is mediated by distinct chromosomal factors. Infect Immun 86: e00141–18.

23) Nashat MA, Luchins KR, Lepherd ML, Riedel ER, Izdebska JN, Lipman NS. 2017. Characterization of Demodex musculi infestation, associated comorbidities, and topographic distribution in a mouse strain with defective adaptive immunity. Comp med 67: 315–329.

24) National Research Council. 2011. Guide for the Care and Use of Laboratory Animals: Eighth Edition. Washington, DC: The National Academies Press.

25) Nigg C, Eaton MD. 1944. Isolation from normal mice of a pneumotropic virus which forms elementary bodies. J Exper Med 79: 497–510.

26) Pal S, Tifra DF, de la Maza LM. 2019. Characterization of the horizontal and vertical sexual transmission of Chlamydia genital infections in a new mouse model. Infect Immun 87: e00834–18.

27) Perry LL, Hughes S. 1999. Chlamydial colonization of multiple mucosae following infection by any mucosal route. Inf Immun 67: 3686–3689.

28) Poston TB, OConnell CM, Girardi J, Sullivan JE, Nagarajan UM, Marinov A, Scurlock AM, Darville T. 2018. T cell-independent gamma interferon and B cells cooperate to prevent mortality associated with disseminated Chlamydia muridarum genital tract infection. Infect Immun. 86: e00143–18.

29) Ramsey KH, Sigar IM, Schripsema JH, Denman CJ, Bowlin AK, Myers GA, Rank RG. 2009. Strain and virulence diversity in the mouse pathogen Chlamydia muridarum. Infect Immun 77: 3284–93.

30) Ramsey KH, Sigar IM, Schripsema JH, Townsend KE, Barry RJ, Peters J, Platt KB. 2015. Detection of Chlamydia infection in Peromyscus species rodents from sylvatic and laboratory sources. Path and Dis 74: ftv129.

31) Rank R. 2007. Chlamydial Diseases, p 326–344. In: Fox JG, Davisson MT, Quimby FW, Barthold SW, Newcomer CE, Smith AL, editors. The mouse in biomedical research, vol 2. Burlington (MA): Elsevier.

32) Rank R, Soderberg LSF, Barron AL. 1985. Chronic Chlamydial genital infection in congenitally athymic nude mice. Infect Immun 48: 847–849.

33) Read TD, Brunham RC, Shen C, Gill SR, Heidelberg JF, White O, Hickey EK, Peerson J, Utterback T, Berry K, Bass S, Linher K, Weidman J, Khouri H, Craven B, Bowman C, Dodson R, Gwinn M, Nelson W, DeBoy R, Kolonay J, McClarty G, Salzberg SL, Eisen J, Fraser CM. 2000. Genome sequences of Chlamydia trachomatis MoPn and Chlamydia pneumoniae AR39. Nucleic Acids Res. 28:1397–1406.

34) Scavizzi F, Bassi C, Lupini L, Guerriero P, Raspa M, Sabbioni S. 2021. A comprehensive approach for microbiota and health monitoring in mouse colonies using metagenomic shotgun sequencing. Anim Microbiome. 3: 53.

35) Starkey MR, Essilfie AT, Horvat JC, Kim RY, Nguyen DH, Beagley KW, Mattes J, Foster PS, Hansbro PM. 2012. Constitutive production of IL-13 promotes early-life Chlamydia respiratory infection and allergic airway disease. Mucosal Immunol 6: 569–579.

36) Stills H. F. Jr, Fox J. G., Paster B. J., Dewhirst F. E. 1991. A “new” Chlamydia sp. strain SFPD isolated from transmissible proliferative ileitis in hamsters. Microbiol Ecol Health Dis 4: S99.

37) Stough JM, Dearth SP, Denny JE, LeCleir GR, Schmidt NW, Campagna SR, Wilhelm SW. 2016. Functional characteristics of the gut microbiome in C57Bl/6 mice differentially susceptible to Plasmodium yoelii. Front Microbiol 7:1520.

38) Su H, Morrison R, Messer R, Whitmire W, Hughes, Caldwell HD. 1999. The effect of doxycycline treatment on the development of protective immunity in a murine model of chlamydial genital infection. J Inf Dis 180: 1252–1258.

39) Sundberg JP, Taylor D, Lorch G, Miller J, Silva KA, Sundberg BA, Roopenian D, Sperling L, Ong D, King LE, Everts H. 2011. Primary follicular dystrophy with scarring dermatitis in C57Bl/6 mouse substrains resembles central centrifugal cicatricial alopecia in humans. Vet Pathol 48: 513–524.

40) Thurlow RW, Arriola R, Soll CE, Lipman NS. 2007. Evaluation of a flash disinfection process for surface decontamination of gamma-irradiated feed packaging. J Am Assoc Lab Anim Sci 46: 46–49.

41) Wooters MA, Kaufhold RM, Field JA, et al. 2009. A real-time quantitative polymerase chain reaction assay for the detection of Chlamydia in the mouse genital tract model. Diagn Micr Infec Dis 63: 140–147.

42) Yeruva L, Spencer N, Bowlin AK, Wang Y, Rank RG. 2013. Chlamydial infection of the gastrointestinal tract: a reservoir for persistent infection. Pathog Dis 68: 89–95.

43) Zhang YX, Fox JG, Ho Y, Zhang L, Stills HF, Smith TF. 1993. Comparison of the major outer-membrane protein (MOMP) gene of mouse pneumonitis (MoPn) and hamster SFPD strains of Chlamydia trachomatis with other Chlamydia strains. Mol Biol Evol 10: 1327–1342.

